# Parental infections disrupt clustered genes encoding related functions required for nervous system development in newborns

**DOI:** 10.1101/448845

**Authors:** Bernard Friedenson

## Abstract

The purpose of this study was to understand the role of infection in the origin of chromosomal anomalies linked to neurodevelopmental disorders. In children with disorders in the development of their nervous systems, chromosome anomalies known to cause these disorders were compared to viruses and bacteria including known teratogens. Results support the explanation that parental infections disrupt elaborate multi-system gene coordination needed for neurodevelopment. Genes essential for neurons, lymphatic drainage, immunity, circulation, angiogenesis, cell barriers, structure, and chromatin activity were all found close together in polyfunctional clusters that were deleted in neurodevelopmental disorders. These deletions account for immune, circulatory, and structural deficits that accompany neurologic deficits. In deleted gene clusters, specific and repetitive human DNA matched infections and passed rigorous artifact tests. In some patients, epigenetic driver mutations were found and may be functionally equivalent to deleting a cluster or changing topologic chromatin interactions because they change access to large chromosome segments. In three families, deleted DNA sequences were associated with intellectual deficits and were not included in any database of genomic variants. These sequences were thousands of bp and unequivocally matched foreign DNAs. Analogous homologies were also found in chromosome anomalies of a recurrent neurodevelopmental disorder. Viral and bacterial DNAs that match repetitive or specific human DNA segments are thus proposed to interfere with highly active break repair during meiosis; sometimes delete polyfunctional clusters, and disable epigenetic drivers. Mis-repaired gametes produce zygotes containing rare chromosome anomalies which cause neurologic disorders and accompanying non-neurologic signs. Neurodevelopmental disorders may be examples of assault on the human genome by foreign DNA with some infections more likely tolerated because they resemble human DNA segments. Further tests of this model await new technology.

**Graphic Abstract:** 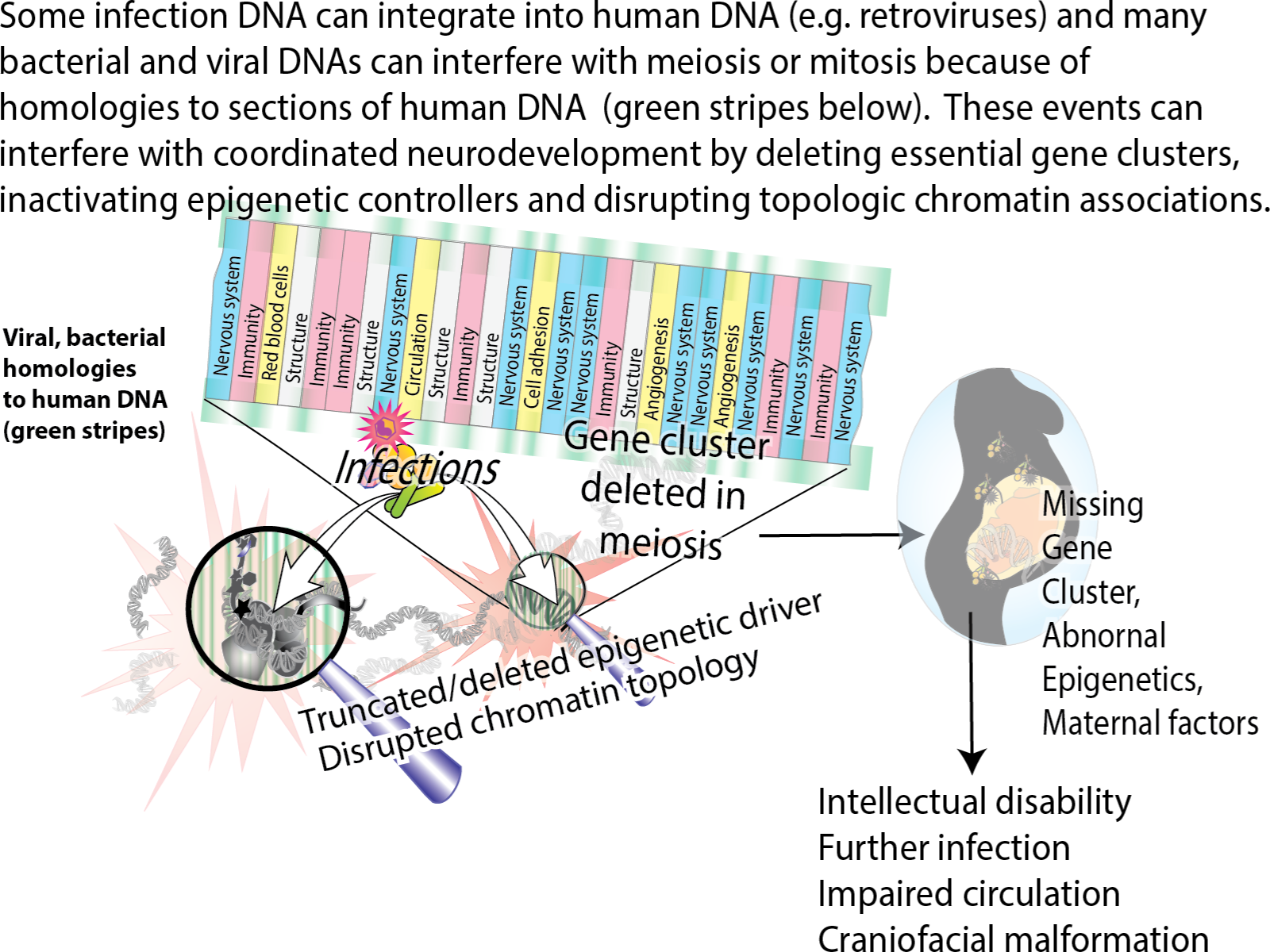

## Introduction

An approach to preventing neurodevelopmental disorders is to gain better understanding of how neurodevelopment is coordinated and then to identify environmental and genomic factors that disrupt it. The development of the nervous system requires tight regulation and coordination of multiple functions essential to protect and nourish neurons. As the nervous system develops, the immune system, the circulatory system, cranial and skeletal systems must all undergo synchronized and coordinated development. Neurodevelopmental disorders follow the disruption of this coordination.

A significant advance in genome sequence level resolution of balanced cytogenetic abnormalities greatly improves the ability to document changes in regulation and dosage for genes critical to the function of the neurologic system. Based on DNA sequence analyses, some chromosome rearrangements have been identified as causing individual congenital disorders because they disrupt genes essential for normal development ^1-3^. There is poor understanding and no effective treatment for many of these overwhelming abnormalities. Signs and symptoms include autism, microcephaly, macrocephaly, behavioral problems, intellectual disability, tantrums, seizures, respiratory problems, spasticity, heart problems, hearing loss, and hallucinations ^1^. Because such abnormalities do not correlate well with prognosis, genetic counseling is difficult and uncertain ^3^.

In congenital neurologic disease, inheritance is usually autosomal dominant and the same chromosomal abnormalities occur in every cell. The genetic events that lead to most neurodevelopmental disorders are not understood ^4^ but several maternal infections and other lifestyle factors are known to interfere. The present work implicates foreign DNA from infections as a source of the chromosome anomalies that cause birth defects. Infections replicate within the CNS by taking advantage of immune deficiencies such as those traced back to deficient microRNA production ^5^ or other gene losses. Disseminated infections can then interfere with the highly active DNA break repair process required during meiosis. Gametes with mis-repaired DNA then cause chromosome anomalies in the zygote. The generation of gametes by meiosis is the most active period of recombination. Double strand breaks initiate meiotic recombination and hundreds of double strand breaks occur ^6^. In contrast to oocytes, meiotic recombination in sperm cells occurs continuously after puberty.

The exact DNA sequences of known pathogenic rearrangements in unique, familial and recurrent congenital disorders ^1-3^ makes it possible to test for the involvement of microbial DNA. Even rare and unique developmental disorders can be screened for homology to infections in the context of altered chromatin structure affecting the immune system, the circulatory system, the formation of the brain-circulation barriers and bone development. This screening may assist counseling, diagnosis, prevention, and early intervention.

The present results show that DNAs in some congenital neurodevelopmental disorders closely matches DNA in multiple infections. The matches extend over long linear stretches of human DNA and often include repetitive human DNA sequences. Neurodevelopmental disorders are proposed to begin when parental infections cause insertions or interfere with meiosis at both repetitive and unique human DNA sequences. The affected sequences are shown to exist as linear clusters of genes closely spaced in two dimensions. Interference from infection can also delete or damage human gene clusters and epigenetic regulators that coordinate neurodevelopment. This microbial interference accounts for immune, circulatory, and structural deficits that accompany neurologic deficits. Congenital neurodevelopmental disorders are thus viewed as resulting from an assault on human DNA by microorganisms and an example of the selection of infecting microorganisms based on their similarity to host DNA.

Generating a viable model based on these results may spur the development of methods for identifying contributions from infections in intractable rare disorders that are not now available. Convergent arguments from testing predictions based on any proposed model might lessen effects of limitations in currently available technology.

## Materials and Methods

### Data Sources

DNA sequences from acquired congenital disorders were from published whole genome sequences at chromosome breakpoints and rearrangement sites ^1-3^. Comparison to multiple databases of microbial sequences determined whether there was significant human homology. Emphasis was on cases with strong evidence that a particular human chromosome rearrangement was pathologic for the congenital disorder.

### Testing for microbial sequences

Hundreds of different private rearrangements in patients with different acquired congenital disorders were tested for homology ^7^ against non-human sequences from microorganisms known to infect humans as follows: Viruses (taxid: 10239), and retroviruses including HIV-1 (taxid:11676), human endogenous retroviruses (Taxids:45617, 87786, 11745, 135201, 166122, 228277and 35268); bacteria (taxid:2); Mycobacteria (taxid:85007); and chlamydias (taxid:51291)

Because homologies represent interspecies similarities, “Discontinuous Megablast” was used. Significant homology (indicated by homology score) occurs when microbial and human DNA sequences have more similarity than expected by chance (E value<=e-10) ^8^. Confirmation of microorganism homologies was done by testing multiple variants of complete microorganism genomes against human genomes.

Various literature analyses have placed Alu repeats into 8 subfamilies having consensus sequences (GenBank; accession numbers U14567 - U14574). Microbial sequences were independently compared to all 8 consensus Alu sequences and to 442 individual AluY sequences.

### Chromosome localizations

The positions of microbial homologies in human chromosomes were determined using BLAT or BLAST. Comparisons were also made to cDNAs based on 107,186 Reference Sequence (RefSeq) RNAs derived from the genome sequence with varying levels of transcript or protein homology support. Tests for contamination by vector sequences in these non-templated sequences were also carried out with the BLAST program. Inserted sequences were also compared to Mus musculus GRCM38.p4 [GCF_000001635.24] chromosomes plus unplaced and unlocalized scaffolds (reference assembly in Annotation Release 106). Homology of inserted sequences to each other was tested using the Needleman and Wunsch algorithm.

## Results

### Interdependent functions are clustered together on the same chromosome segment

The nervous system shows a close relationship to structures essential for immunity, circulation, cell barriers and protective enclosures. In chromosome segments deleted in neurologic disorders, genes essential for all these functions must develop in concert. Genes for these related functions are located close to each other on the same linear segment of a chromosome (Fig. 1). Deletions at 4q34 in patient DGAP161 and at 19q12-13.11 in patient DGAP125 are shown in Fig. 1, as examples that are representative of the other large chromosomal deletions. The genes within each deletion in each of four categories tested are color coded in the figure. Deletion of these clustered arrangements has been correlated with serious neurodevelopmental disorders^1^. Alternatively-spliced forms of the same gene may encode for different functions that must be synchronized and coordinated among diverse cells. Multiple functions in different cell types are commonly found. Hormonal signaling represents a major control mechanism^9^.

**Fig. 1.**
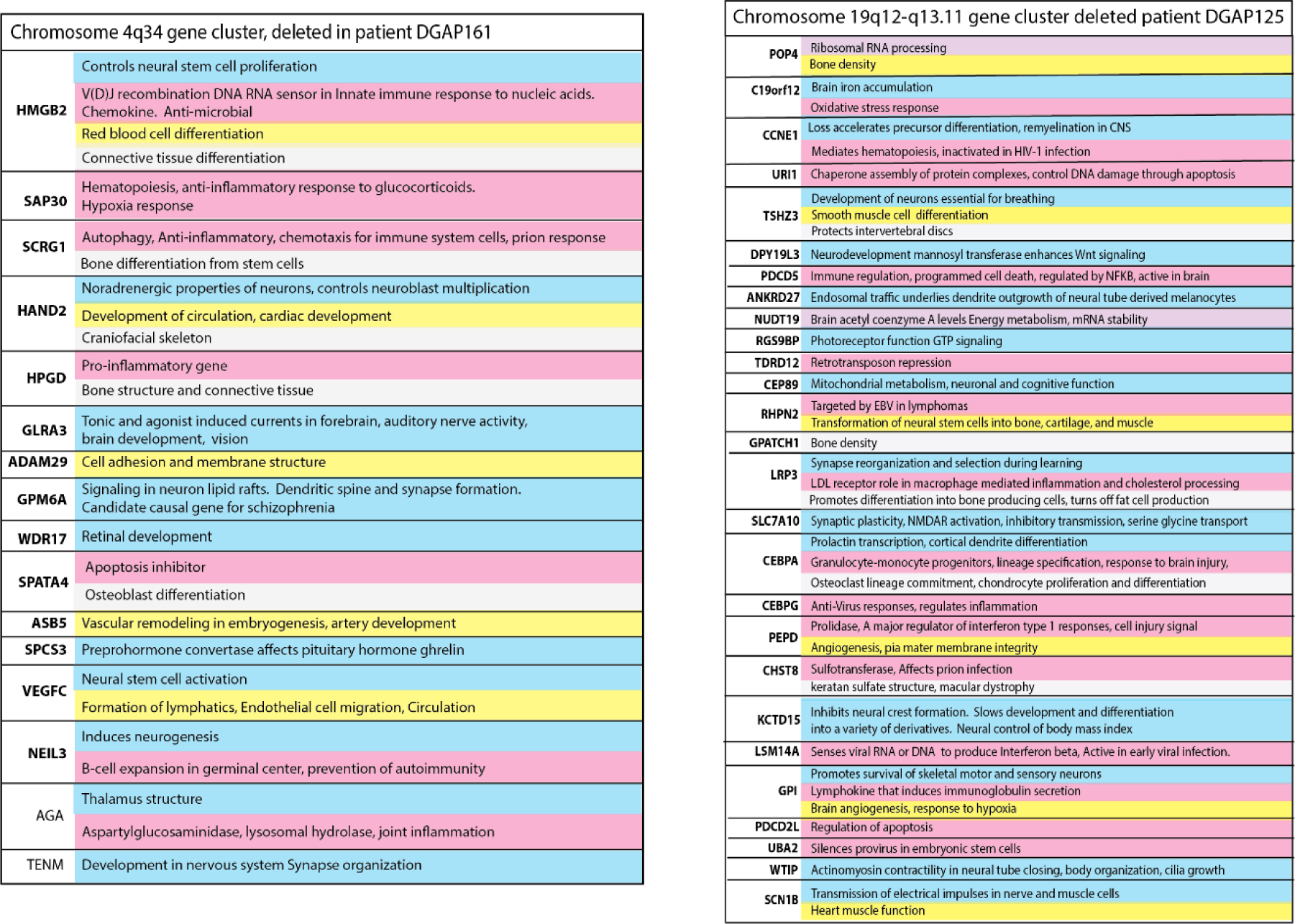
Close relationships between the nervous system and other essential developmental functions. Genes for multiple interdependent systems appear in clusters on the same chromosomes. Nervous system genes are near genes essential for the immune system, connections to lymphatic circulation, ability to form tight junctions, structural enclosures and chromatin control. Clusters of genes encoding these and other interdependent functions on chromosome segments are deleted in private neurodevelopmental disorders. These losses increase susceptibility to infections which are homologous to long stretches of human DNA. In the two examples shown, genes are listed in the order they occur on two typical chromosome deletions. Functions are to the right of the gene symbol. Blue genes are associated with the nervous system, pink with the immune system. Yellow genes have functions associated with angiogenesis, lymphangiogenesis or cell barriers. Genes for development of essential bone structure or connective tissues needed to protect and house the nervous system are light grey. Genes colored purple code for general functions required by all cells. The same gene may function differently in different cellular locations.

### Many deleted gene clusters include long stretches of DNA strongly related to infections

To investigate the chromosomal segment deletions that cause neurodevelopmental disorders, homologies to infection were tested in sequences within and flanking deleted clusters. Strong homologies to infections were interspersed. To demonstrate the extent of these relationships, deleted 4q34 chromosome segment (Patient DGAP161 ^1^) was tested for homology to microbes in 200 kb chunks. Fig. 2 shows that stretches of homology to microorganisms are distributed throughout chromosome 4q34. Only 10 homologous microbes are shown for each 200 kb division, but there are up to hundreds, giving a total of many thousands of potentially homologous microorganisms throughout the 4q34 deletion.

**Fig. 2.**
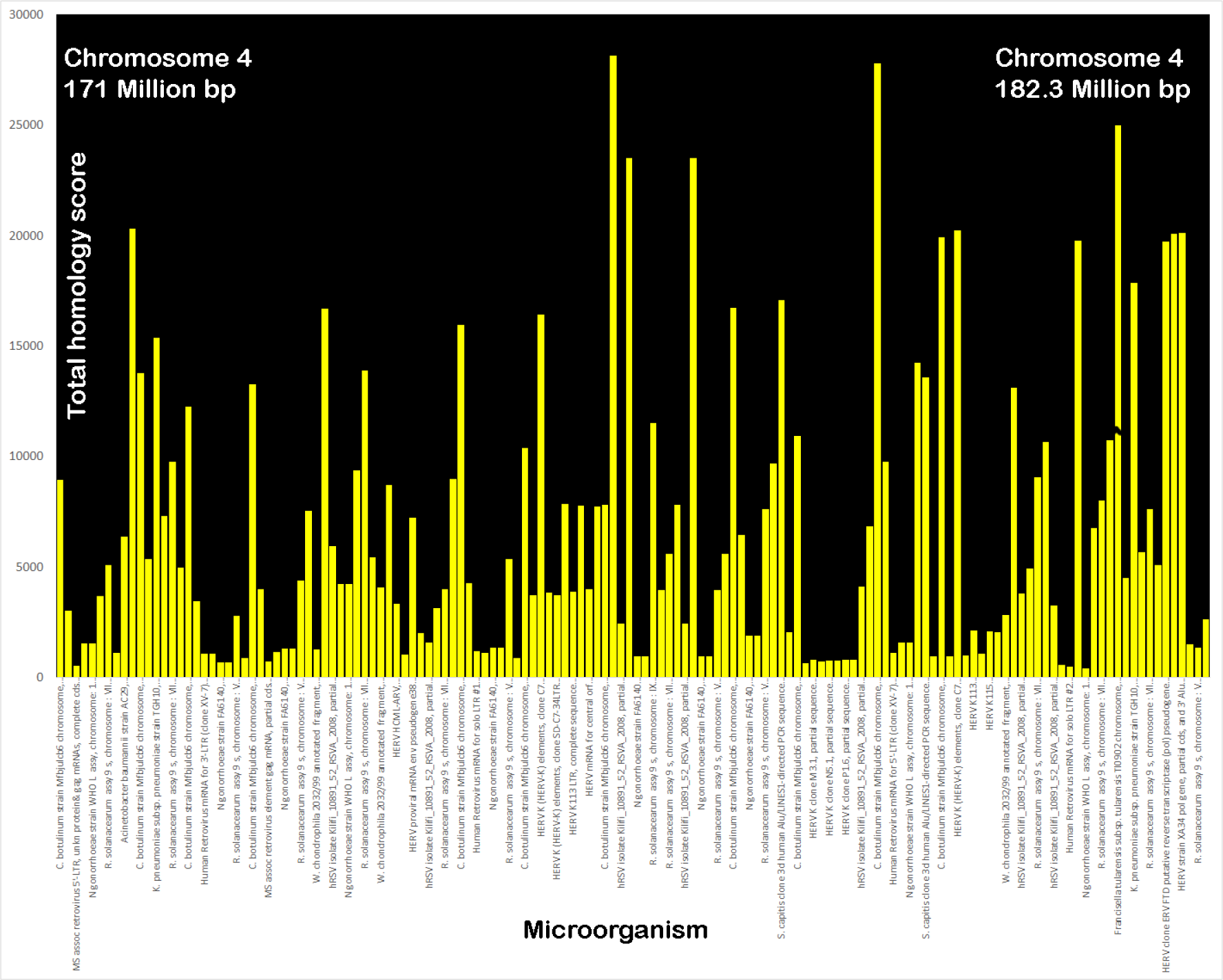
Example of total homologies to microbial sequences dispersed throughout 4q34 deleted chromosome segment in patient DGAP161. The top 10 homologies are shown for each 200,000 bp segment and are listed within each of these 57 segments in order of their scores. The Y axis plots the total homology scores and the x axis lists the individual microorganism. The mean E value was 5.3e-90 (range 0.0 to 1e-87). F. tularensis had values of 33,368 and 79,059 which were truncated at 25000 to show more detail for the other microbial matches.

A deeper analysis of the functions of genes within deleted chromosome segments predicts a predisposition to infection and correlates with symptoms of the developmental disorder. For example, deletion of chromosome 19q12-q13.11 occurs in patient DGAP125 (Fig. 1). The deletion causes deficits in silencing provirus, repressing retrotransposons, responding to viruses, specifying immune cell lineages, and regulating apoptosis. The deleted 19q12-q13.11 band in patient DGAP125 had 76 matches to *clostridium botulinum* strain Mfbjuicb6. Many of these matches extended over 2800 bp at 72-73% homology (E = 0.0). There were also strong matches to Waddlia chondrophila and N. gonorrhoeae. Other shorter chromosomal anomalies are present in patient DGAP125. Stealth and HIV-1 teratogens and multiple other infections are homologous to these shorter anomalies (Fig. 3).

**Fig. 3.**
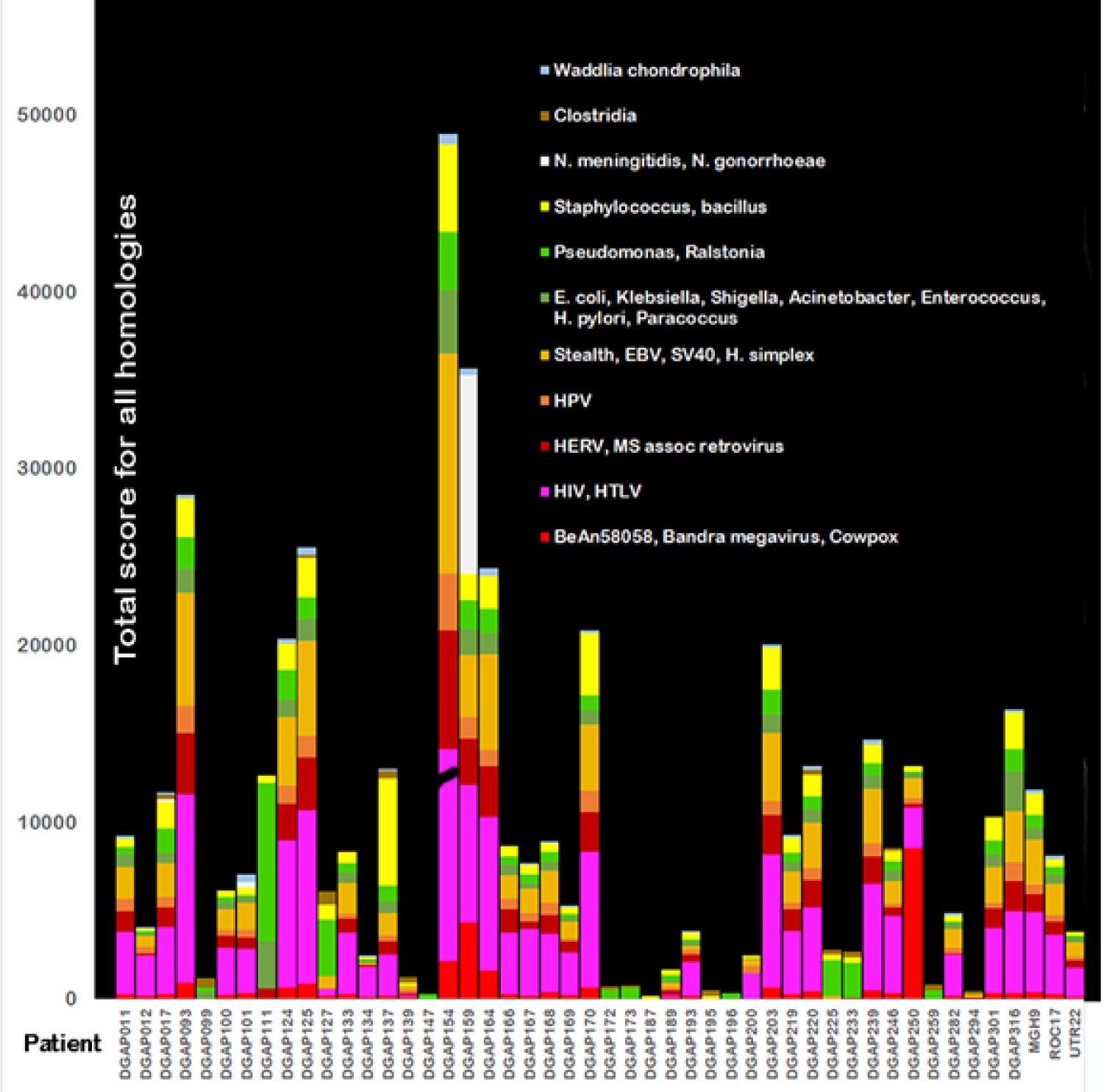
Total scores for homologies among microorganisms and human chromosomal anomalies in 48 patients with neurodevelopmental disorders. The contributions from all homology scores for groupings or individual microorganisms that match the chromosomal anomaly in each patient are added together to form each bar in the graph. The microorganism homology scores are color-coded according to the type of individual microorganism. Of 1986 E values, the mean was 7.3e-13 [Range 8e-11 to 2E-168]. Some of the microorganisms found are not normally pathogenic but can become opportunistic pathogens if the immune system is impaired. Endogenous retroviral sequences may interfere with recombination or repair within a single cell. Total homology between DGAP154 and HIV/HTLV (24,873) was truncated at 12,000 to prevent obscuring other data.

The large deletion in 19q12-q13.11 not only makes patient DGAP125 especially susceptible to infections but also deletes coordinated genes that control breathing, synapse organization and plasticity, neural crest formation, transmission of nerve impulses; development of brain circulation; and production of connective tissue and bone. Other chromosome deletions with clustered genes have deficits in neuronal accessory functions like those in DGAP125.

### Microbial DNA homologies in areas around a mutated epigenetic driver gene

Gene changes identified as underlying phenotypic drivers of congenital neurologic diseases ^1^ were tested against published evidence for their ability to act as epigenetic regulators and effectors of functions like those found in the chromosomal deletions. Table 1 shows that most pathogenic driver genes are epigenetic regulators.

**Table 1.** Epigenetic functions of mutated and deleted genes in neurodevelopmental disorders relate neurologic deficits to deficits in the immune system, the circulatory system and structural genes.

| Patient | Proposed primary phenotypic drivers of anomaly or disrupted genes. Genes in bold type are related to epigenetic control | Genes with pathogenic neurologic mutation <sup>11</sup> associated with nervous system function | Genes deleted or mutated linked to immunity, infection | Genes deleted or mutated with linked to circulation and blood brain barrier | Genes deleted or mutated linked to bone, structural requirements |
| --- | --- | --- | --- | --- | --- |
| DGAP002 | 14q12-q21.1 deleted. **FOXAI** in the deletion mediates chromatin remodeling | NOVA1, AKAP6 | Fox G1, NFKB1A, NKX2, BAZ1A, FoxA1 | FoxA1 | FoxA1 |
| DGAP011 | **FGFR1** Receptor for gene that regulates chromatin organization during development | FGFR1 | FGFR1 | FGFR1 | FGFR1 |
| DGAP012 | **PHF21A**, chromatin regulator disrupted at breakpoint, encodes part of histone deacetylase complex | Encodes BHC80, BRAF35 in histone deacetylase complex. | Histone deacetylase | Histone deacetylase. | Not found |
| DGAP093 | **CDKL5** gene disrupted at breakpoint, phosphorylates HDAC4 histone deacetylase causing cytoplasmic retention <sup>12</sup>. | CDKL5 Mediates MECP2 phosphorylation. . | CDKL5 | CDKL5 | Not found |
| DGAP096 | **WAC** at breakpoint, disrupted by translocation regulates histone ubiquitination | WAC. | WAC | WAC | Not found |
| DGAP099 | **ZBTB20** binds to genes that control chromatin architecture including MEF2c and SAT2b | ZBTB20 | ZBTB20 | ZBTB20 | ZBTB20 |
| DGAP100 | **KDM6** Histone Demethylase and Epigenetic switch to specify stem cell fate <sup>13</sup> | KDM6A | KDM6A | KDM6A | KDM6A |
| DGAP112 | 12p12.1-p11.22 deleted. **SOX5**, a histone demethylase at breakpoint is disrupted by rearrangement. MED21 at breakpoint may be required to displace histones to enable heat-shock response <sup>14</sup>. | SOX5 | SOX5, KRAS, MED 21 | SOX5, KRAS | SOX5 |
| DGAP124 | **NRXN1**, repressive histone marker at rearrangement breakpoint, controls choice of NRXN splicing isoform. Repressed by histone methyl-transferase <sup>15</sup> | NRXN1 | NRXN1 | NRXN1 | Not found |
| DGAP127 | **PAK3**, compensates for PAK1 which interacts with histone H3 | PAK3 | PAK3 | PAK3 | Not found |
| DGAP133 | 6q13-q14.1 deleted includes **HMGN3** chromatin binding protein that stimulates acetylation <sup>16</sup> | SMAP1, B3GAT2, OGFR1, RIMS1, SLC17A5, FILIP1, LCA5, SENP6, etc (See Fig 1) | KHDC1, EEFI1A, MYO6, IRAK1BP1, B3GAT2, etc. (see Fig 1) | Collagen genes 19A, 12A, COX7A2 | Col9A |
| DGAP139 | 13q14.2 deleted Includes **SETDB2** <sup>17</sup> (See Fig. 1) | SUCLA2, ITM2B, RCBTB2, CYSLTR2, KPNA3 | RBI, NUDT15, DLEU1,2, PHF11, SETDB2, KCNRG | ITM2B, CYSLTR2, LPAR6, DLEU2 | DLeu2 |
| DGAP142 | **MBD5** Epigenetic regulator Histone acetylation disrupted by rearrangement. | MBD5 | MBD5 | MBD5 | Not found |
| DGAPI45 | **EFTUD2** [RNA splicing regulator in spliceosome] at rearrangement breakpoint. Connects methylated histone to RNA splicing<sup>18</sup> | EFTUD2 | EFTUD2 | EFTUD2 | EFTUD2 |
| DGAPI47 | **NALCN** Disrupted at breakpoint | NALCN | NALCN | Not found | Not found |
| DGAPI54 | Xq25 (Duplication) **XIAP, STAG2**<sup>19</sup> | XIAP, THOC2, GRIA3 | XIAP | SMAD5 | SMAD5 |
| DGAPI55 | **EHMT1**/GLP: Encodes a histone methyltransferase, an epigenetic regulator. | EHMT1 | EHMT1 | EHMT1 | EHMT1 |
| DGAPI57 | **FOXPI** "PRMT5 recruitment to the FOXPI promoter facilitates H3R2me2s, SET1 recruitment, H3K4me3, and gene expression"<sup>20</sup> Disrupted by rearrangement. | FOXPI | FOXPI | FOXPI | FOXPI |
| DGAPI59 | 10p15.3-p14 Deletion includes **GATA3** See Fig. 1. GATA3 controls enhancer accessibility, recruits chromatin remodeler to reprogram cells. | GATA3 | GATA3 | GATA3 | GATA3 |
| DGAPI61 | 4q34 (See Fig. 1) **HMGB2**, a chromatin protein affecting epigenetic status<sup>21</sup> | AGA, SPCS3, TENM | HMGB2, HPGD, VEGFc ADAM29 | VEGFc, ADAM29 | HMGB2, VEGFc |
| DGAPI64 | **NODAL/activin a** targets KDM6B (epigenetic regulator of neuronal plasticity) signaling in mouse brain. Rearrangement also disrupts **TET1** (a methylation eraser (demethylase)) | NODAL | NODAL | NODAL | NODAL |
| DGAPI66 | **SCN1A** | SCN1A | SCN1A | Not found | Not found |
| DGAPI69 | **NR2F1** disrupted by rearrangement, Induces global chromatin repression | NR2F1 | Not found | NR2F1 | NR2F1 |
| DGAPI73 | 11p14.2 **FIBIN, BBOX1, SLC5A12, ANO3** | ANO3, BBOX1 | Mucin15 | Not found | Not found |
| DGAPI86 | **NR5A1** disrupted, participates in resetting pluripotency | NR5A1 | NR5A1 | NR5A1 | Not found |
| DGAPI89 | **SOX5** is a histone demethylase | SOX5 | SOX5 | SOX5 | SOX5 |
| DGAPI90 | **SMS** Drives interactions among nucleosomes. | Spermine synthetase gene, spermine. | Spermine synthetase gene, spermine. | Spermine synthetase gene, spermine. | Spermine synthetase gene, spermine. |
| DGAPI93 | **SPAST** | SPAST | SPAST | SPAST | SPAST |
| DGAPI99 | **NOTCH2**, disrupted at breakpoint by rearrangement. | NOTCH2 | NOTCH2 | NOTCH2 | NOTCH2 |
| DGAP201 | **AUTS2** Disrupted at breakpoint by rearrangement. Associated with chromatin structure | AUTS2 | AUTS2 | Not found | Not found |
| DGAP202 | **KDM6A** Histone di/tri demethylase disrupted at breakpoint by rearrangement. | KDM6A | KDM6a | KDM6a | KDM6a |
| DGAP211 | **SATB2** Disrupted at breakpoint by rearrangement. Epigenetic functions, interacts with chromatin remodelers. | SATB2 | SATB2 | Not found | SATB2 |
| DGAP219 | **CUL3** Ubiquitin ligase recruited to regulate chromatin pattern of lymphoid effector programs. Disrupted at breakpoint of rearrangement. | CUL3 | CUL3 | CUL3 | CUL3 |
| DGAP232 | **SNRPN-SNURF** / SMN; PWCR; SM-D; sm-N; RT-LI; HCERN3; SNRNP-N; Required for epigenetic imprinting. | SNRPN-SNURF | SNRPN-SNURF | Not found | SNRPN-SNURF |
| DGAP235 | **MBD5** Epigenetic regulator histone acetylation disrupted at breakpoint of rearrangement. | MBD5 | MBD5 | MBD5 | Not found |
| DGAP239 | **CHD7** Chromatin remodeler. | CHD7 | CHD7 | Not found | CHD7 |
| DGAP244 | **CTNND2** | CTNND2 | CTNND2 | Not found | CTNND2 |
| DGAP278 | **SNRPN-SNURF** / SMN; PWCR; SM-D; sm-N; RT-LI; HCERN3; SNRNP-N; Required for epigenetic imprinting | SNRPN-SNURF | SNRPN-SNURF | Not found | SNRPN-SNURF |
| DGAP301 | **MEF2C** , Associated with disease histone hypermethylation<sup>22</sup>.. Disrupted by rearrangement | MEF2C | MEF2C | MEF2C | MEF2C |
| DGAP316 | 18p11.32-p11.22 deletion: **PHF21A** PHF21A Encodes BHC80, a component of a BRAF35 histone deacetylase complex. | PHF21A | PHF21A | YES1, MYOM1, EPB41L3, ARHGAP28, LAMA1, PTPRM, MTCL1, TWSG1, RALBP1, RAB31 | SMCHD1, TGIF1, LAMA1, PTPRM, TWSGH1, |
| DGAP317 | 6q14.1 TBX18: IRAK1BP1; IBTK | LCA5, ELOV4, HTR1B, DOPEY | IRAK1BP1, TBX18, IBTK, NFKB | COX7A | Not found |
| MGH7 | **GRIN2B** Reflects changes in methylation levels <sup>23</sup> | GRIN2B | GRIN2B | Not found | Not found |
| MGH8 | **CHD8**, chromatin insulator blocks effects of nearby enhancers. CHD8 interacts with the insulator binding protein CTCF to insulate and epigenetically regulate <sup>24</sup>. | CHD8 | CHD8 | Not found | Not found |
| MGH9 | **TCF4** regulated by histone deacetylases. Binding sites overlap histone acetylation of enhancers <sup>25</sup> | TCF4 | TCF4 | TCF4 | TCF4 |
| NIJ2 | **PHIP**, linked to histone methylation <sup>26</sup> Disrupted at breakpoint by rearrangement. | PHIP, MYO6 | PHIP, MYO6 | PHIP, MYO6 | PHIP, MYO6 |
| NIJ5 | *ILIRAPL1* | ILIRAPL1 | ILIRAPL1 | Not found | Skeletal growth |
| NIJ6 | **KAT6B** / MORF MYST4 Lysine acetyl transferase. | KAT6B | KAT6B | KAT6B | KAT6B |
| NIJ14 | **NFIA**, regulated by chromatin structure <sup>27</sup> | NFIA | NFIA | NFIA | NFIA |
| NIJ15, NIJ6 | *MYT1L* | MYT1L | MYT1L | Not found | Not found |
| ROC4 | *CAMTA1* | CAMTA1 | CAMTA1 | CAMTA1 | CAMTA1 |
| ROC17 | **AUTS2** Associated with H3K4me3 and epigenetic control | AUTS2 | AUTS2 | Not found | Not found |
| ROC23 | *TCF12* | TCF12 | TCF12 | TCF12 | TCF12 |
| ROC43 | **SOX5** is a histone demethylase | SOX5 | SOX5 | SOX5 | SOX5 |
| ROC62 | **SNRPN-SNURF** Required for epigenetic imprinting | SNRPN-SNURF | SNRPN-SNURF | Not found | SNRPN-SNURF |
| UTR7 | **NFIX**, controlled by methylation | NFIX | NFIX | NFIX | NFIX |
| UTR12 | **DYRK1A**, histone phosphorylation | DYRK1A | DYRK1A | DYRK1A | DYRK1A |
| UTR13 | **MBD5**, Epigenetic regulator disrupted in topology associated domain | MBD5 | MBD5 | MBD5 | Not found |
| UTR17 | **ZBTB20**, binds to genes that control chromatin architecture including MEF2c and SAT2b) | ZBTB20 | ZBTB20 | ZBTB20 | ZBTB20 |
| UTR20 | *FOXP2* | FOXP2 | FOXP2 | FOXP2 | FOXP2 |
| UTR21 | **NSD1** lysine methylation | NSD1 | NSD1 | NSD1 | NSD1 |
| UTR22 | 2q24.3 SCN9A | SCN9A, TTC21B, STK39 | NOSTRIN, CERS6 | FIGN, XIRP2, CERS6 | TTC21B |
| Affected member of family 1 | **NCALD1** exists in myristoyl and non myristoyl forms. Histone deacetylase 11 is a myristoyl hydrolase <sup>28</sup>. | NCALD1 | NCALD1 | Not found | NCALD1 |
| Affected member of family 1 | *NPL* - N-acetylneuraminatase lyases regulate cellular sialic acid concentrations by mediating its reversible conversion into N-acetylmannosamine and pyruvate | NPL | NPL | NPL | NPL |
| Affected member of family 1 | **BCAR3**. A candidate epigenetic mediator for increased adult body mass index in socially disadvantaged females <sup>29</sup> | Not found | Not found | BCAR3 | BCAR3 |
| Affected member of family 2 | **ZNF423** - epigenetic modifications associated with obesity <sup>30</sup>, methylation marker for age prediction. | ZNF423 | ZNF423 |  | ZNF423 |
| Affected member of family 3 | **RAP1GDS1** - **RAP1** GTP-GDP dissociation stimulator <sup>31</sup>. Antagonizes some histones. | RAP1GDS1 - RAP1 | RAP1GDS1 - RAP1 | RAP1GDS1 - RAP1 | RAP1GDS1 - RAP1 |

Fig. 3 graphically relates DNA sequences in all viruses and bacteria in the NCBI database to DNA sequences around breakage sites based on outstanding data published by the study of Redin and colleagues ^1^. There may also be cryptic rearrangements elsewhere ^2,3,10^. The figure shows wide differences in the total homology scores of microbes vs. humans in individual disorders. Patient DGAP154 has total homology scores of well over 50,000 while other patients (DGAP142, 172, and 173) are well below 1000.

### Pathogenic driver gene mutations are roughly equivalent to large chromosome deletions

Because most identified driver genes of neurodevelopmental disorders ^1^ are epigenetic regulators or effectors (Table 1), the functions they control in individual patients and in families with members affected by neurodevelopmental disorders ^3^ were compared to genes in pathogenic chromosomal deletions. Like clustered chromosomal deletions, virtually all pathogenic driver genes have strong effects on the immune system, angiogenesis, circulation and craniofacial development. Fig. 4 summarizes how the functions of damaged epigenetic drivers are distributed.

**Fig. 4.**
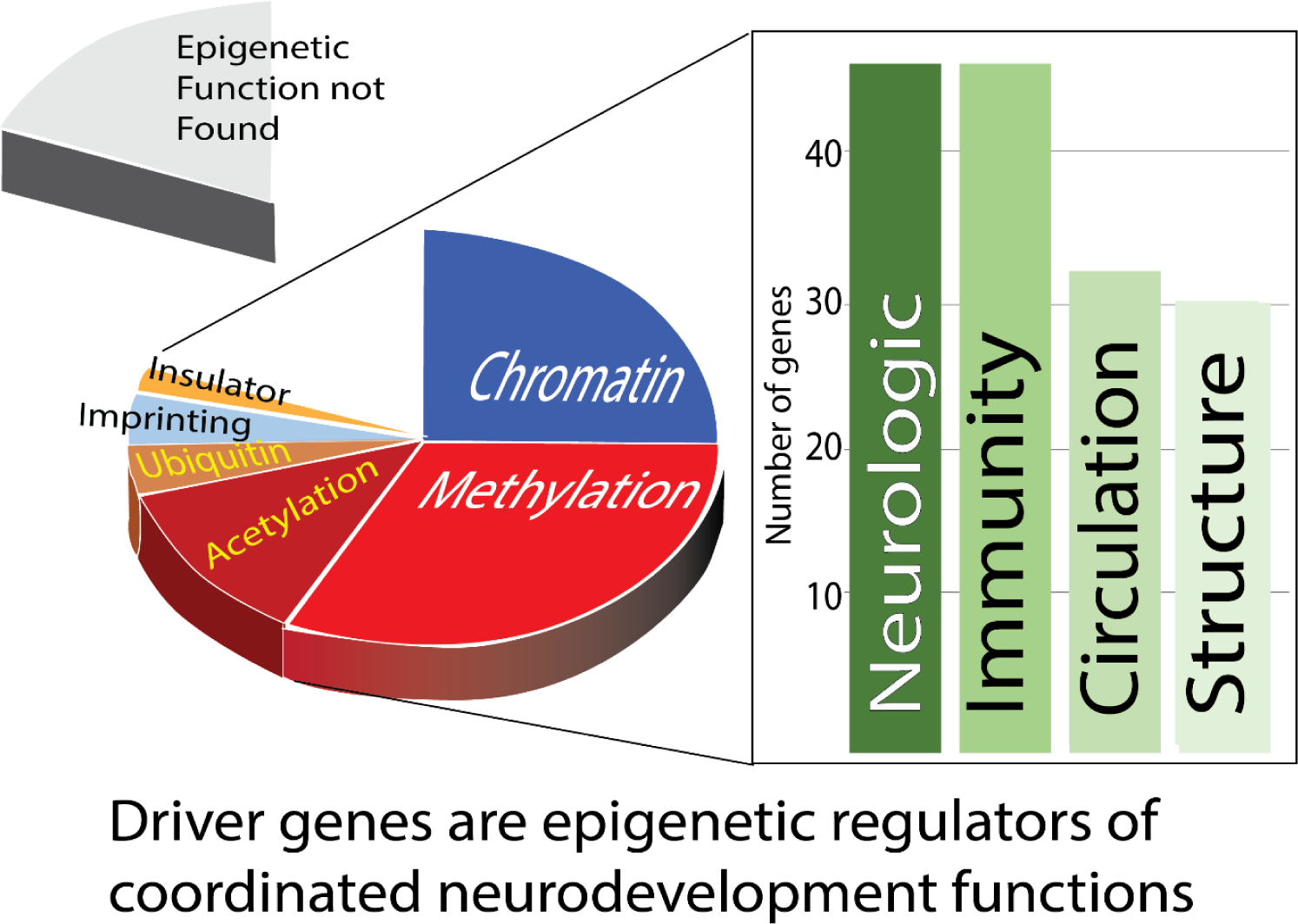
Driver genes truncated or deleted in congenital neurodevelopmental disorders are mainly epigenetic regulators or effectors. The pie chart shows the percentages of 46 driver genes with indicated epigenetic functions. Loss of these driver gene functions then impacts a group of functions that must be synchronized during the complex process of neurodevelopment. These are the same general functions lost in deleted gene clusters.

### Multiple infections identified by homology match signs and symptoms of neurodevelopmental disorders

In 48 patients ^1^ shown in Fig. 3, multiple infections are candidates to cause the signs and symptoms in each individual (Table 2). 8 patients have growth retardation or “short stature” which can be caused by exposure to many infections in the second or third trimester. The absence, delay, or impairment of speech were found in 20 of 48 patients ^1^. Multiple infections can cause these problems. For instance, HIV-1 causes white matter lesions associated with language impairments and impaired fetal growth. There are nearly 50 matches to HIV-1 DNA in the chromosome anomalies of 35 patients.

**Table 2.** Recurrent infections found to have homology to chromosomal abnormalities in neurologic birth defects can cause developmental defects.

| ***Recurrent Infections matching DNA rearrangements in congenital neurologic disorders*** | ***CNS, physiologic and or teratogenic effects if known.*** | ***Numbers of Patients with chromosomal abnormalities with significant homologies to this infection*** |
| --- | --- | --- |
| **HIV-1, HIV-2** | Impaired fetal growth, premature delivery, chorioamnionitis, deciduitis, immunodeficiency. HIV-1 causes white matter lesions associated with language impairments. HIV-1 infects CNS by targeting microglial cells. | 600 matches in 36 patients |
| **HPV16** | Part of a group of infections that coexist with other viral infections, bacterial infections, and chemicals in autism syndrome disorders <sup>32</sup>. | 35 patients |
| **Staphylococcal infection, (s. aureus, s. capitis, s. epidermidis)** | May infect brain and interfere with normal nerve transmission. Causes meningitis, Brain abscess. Major hospital pathogens | 206 matches in 40 patients |
| **Stealth viruses, Stealth virus I, CMV, HHV-4/EBV, herpes simplex, SV40** | Associated with neurologic impairments, hearing loss, ophthalmic problems. | 425 matches in 39 patients |
| **Ralstonia solanacearum** | Opportunistic bacterial pathogen, associated with pneumonia and neonatal sepsis, especially in patients with compromised immunity on ventilator. Has some resemblance to other human pathogens including Borrelia, Bordetella, Burkholderia, and a "blood disease pathogen." | 158 matches in 44 patients |
| **HTLV-I** | Associated with neuropathy. Immune mediated disease of the nervous system affects spinal cord and peripheral nerves. Fetal neural cells are highly susceptible to infection | 24 patients |
| **BeAn 58058 virus, Bandra megavirus** | BeAn 58058 is almost identical (97%) to cotia virus which can infect human cells <sup>33</sup>. Represents a distinct branch of poxviruses <sup>34</sup> | 50 matches in 36 patients |
| **Klebsiella pneumoniae, e coli** | Neonatal pneumonia and sepsis. | 35 patients |
| **Human respiratory syncytial virus Kilifi** | Neonatal pneumonia, trouble breathing | 7 patients: DGAP099, DGAPI27, DGAPI37, DGHAPI59, DGAPI67, DGAPI72, DGAP225 |
| **Waddlia Chondrophila** | Recently discovered. Causes preterm birth, miscarriage, spontaneous abortion. Chlamydia like micro-organism. Survives in macrophages, multiplies in endometrial cells | 26 different patients |
| **Neisseria meningitidis, N. gonorrhoeae** | Bacteria that can cause damage to human DNA. Premature birth, miscarriage, severe eye infection in infant. Antibodies to N. gonorrhoeae impair out-growth of neurites, and cross react with specific brain proteins. | Patient DGAPI59 shows 25 different homologies to N. meningitidis. DGAP017 and in DGAPI01 have homology to N. gonorrhoeae. |
| ***Pseudomonas putida*** | Gram-negative bacterium found in cases of acute bacterial meningitis <sup>35</sup>. Reported in contaminated water and heparinized flush solutions <sup>36</sup> | 13 patients: DGAP012, DGAP93, DGAP100, DGAP125, DGAP133, DGAP134, DGAP154, DGAP159, DGAP167, DGAP169, DGAP170, DGAP220, ROC17 |
| ***Exiguobacterium oxidotolerans*** | High catalase activity may disable immune defenses depending on peroxide. | DGAP127, DGAP225 |

Stealth viruses are frequent matches to chromosome anomalies with about 35 matching sequences. Stealth viruses are mostly derivatives of herpes viruses including cytomegalovirus (CMV) that emerge in immunosuppressed patients or have other mechanisms to avoid recognition by the immune system. Stealth virus 1 (Table 2) is Simian CMV (strain Colburn) African green monkey DMV, but closely related viruses with up to 95% sequence identity have been isolated from human patients. First trimester CMV infection can cause severe cerebral abnormalities followed by neurologic symptoms ^37^. CMV is also a common cause of congenital deafness and can cause visual abnormalities. 27 of the 48 neurodevelopmental patients had hearing loss or deafness. Herpes simplex virus is another stealth virus that directly infects the central nervous system. Herpes simplex can cause seizures (reported for 9 patients).

Patient DGAP159 has the strongest homology to N. meningitidis (Fig. 3). N. meningitidis causes bacterial meningitis and is a known cause of neurodevelopmental defects. Signs and symptoms in patient DGAP159 are consistent with residual effects of bacterial meningitis including hearing loss, developmental delay, speech failure and visual problems.

### DNA from microbes not classified as teratogens can also affect fetal development

Not only known teratogens, but almost all microbial DNA matching human chromosome anomalies in congenital neurologic disorders can have profound effects on a fetus or neonate (Table 2). Fig. 3 implies that any source of foreign DNA may contribute to developmental errors. Multiple matches represent homologies to human DNA among stretches of viral and bacterial DNA. Homologies found appear independent of whether microbes can insert into human DNA. HIV, HPV16, HTLV-1, *k. pneumoniae, ralstonia solanacearum, s. aureus,* and stealth viruses are all found in at least 10 patients. 38 of 48 rearrangements include sequences that match one or more viruses, and 44 match bacterial sequences. In most chromosomal anomalies, multiple infections match a given anomalous DNA sequence. It is likely that most or all of these infections can interfere with neurodevelopment (Tables 1-2). Fig. 3 includes endogenous retroviral sequences, mostly represent crippled infections that cannot travel between cells because they lack *env* genes needed to exit infected cells. However, HERV sequences are numerous and as retrotransposons, they can still interfere with recombination and DNA repair within their home cell.

### Deleted segments in familial chromosome anomalies point toward a general mechanism for infection as a cause of neurodevelopmental disorders

Effects of balanced translocations segregate only somewhat with the presence of neurodevelopmental disorders. These discrepancies are not extremely common but they emphasize the need to consider at least some genomes in three-dimensions ^38^ and to consider cryptic anomalies. Genome sequencing of the entire family may be necessary ^3^ because members of some families carry balanced chromosomal translocations but do not have signs and symptoms of neurodevelopmental disease. In three of four families with familial balanced chromosomal translocations, patient specific unbalanced deletions were also found but the results did not overlap any database of human reference genomes ^3^. Unbalanced deletions in the three families were strongly homologous to infections or other foreign DNAs (Fig. 5). In agreement with results in Table 1, critical genes linked to epigenetic modifications were among those interrupted by cryptic rearrangements.

**Fig. 5.**
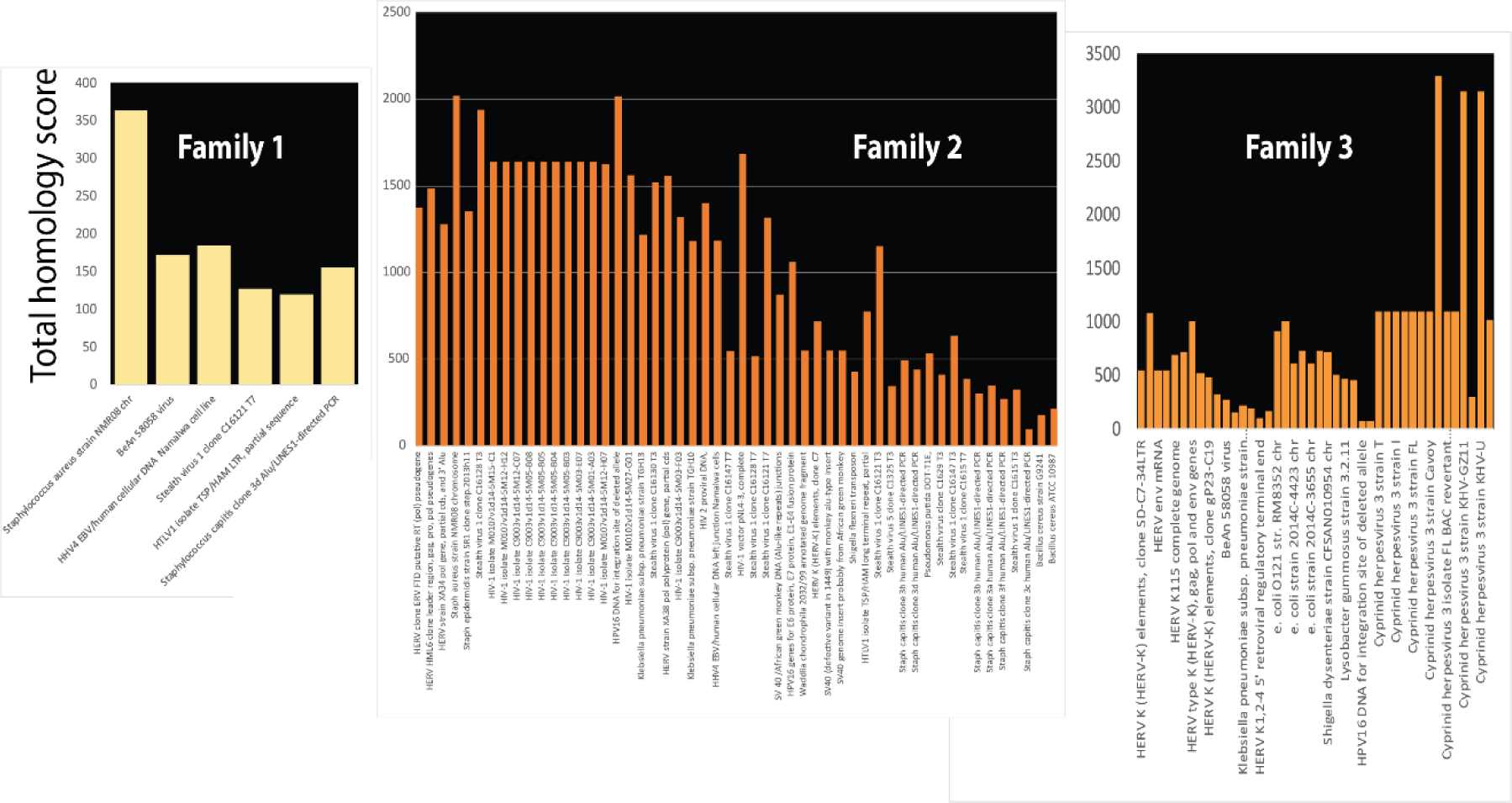
Unbalanced deletions in three families with some affected members are homologous to microorganisms. Individuals affected by intellectual disability from families with apparently balanced translocations have other chromosome abnormalities including unbalanced deletions, insertions ^3^. Chromosome deletions in each family are shown as homologous to bacterial and viral DNA, but insertions were also found to have similar homologies. Total homology scores for bacterial and viral homologies in each of the three affected patients are plotted.

### Chromosome regions with microbial homologies in a recurrent translocation are only deleted in affected family members

Recurrent de novo translocations between chromosomes 11 and 22 have so far only been detected during spermatogenesis and have been attributed to palindromic structures that induce genomic instability ^2^. The recurrent breakpoint t(8;22)(q24.13;q11.21) ^2^ was tested to determine whether palindromic effects might arise because microbial infection interferes with normal chromatin structures. Fig. 6 shows strong homology to bacterial and viral sequences in the recurrent breakpoint region. Sequences from an unaffected mother carry a balanced translocation rearrangement ^2^ with homologies to more diverse microbes than are present in affected cases (Fig. 6). Parts of a chromosome segment with homologies to infection have been deleted in affected family members but not in unaffected ones.

**Fig. 6.**
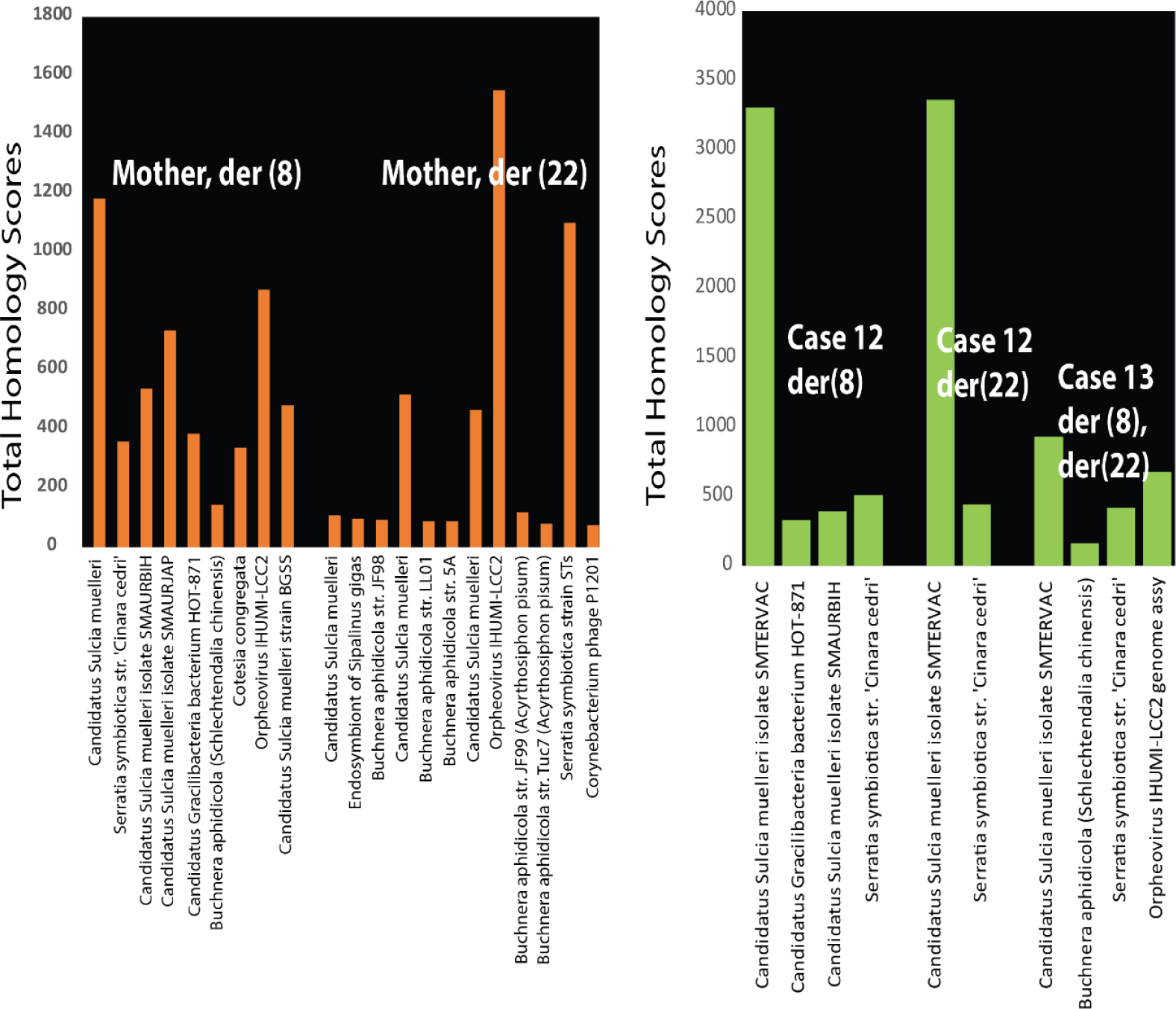
Microbial homologies in cases from a maternal recurrent translocation. Parental translocation t(8;22) in Family FHU13-027 from Mishra and colleagues ^2^ is homologous to microorganisms as shown for der(8) (left) and for der(22) (right). The sequence is from a normal healthy mother who inherited from a female proband the balanced translocation t(8;22). More limited sets of micro-organisms are present in affected case 12 and case 13 (green bars) implying that DNA homologous to the other microorganisms has been lost.

### A model for infection interference in neurodevelopment

Infection DNA occurs in neurodevelopmental disorders that undergo autosomal dominant inheritance suggesting microbial DNA interferes with gamete generation in one parent. The mechanism proposed in Fig. 7 is based on significant homologies with microbial infections on multiple human chromosomes. Large amounts of foreign infection DNA present during human meiosis with its many double strand breaks and the most active period of recombination produce defective gametes. Errors in spermatogenesis underlie a prevalent and recurrent gene rearrangement that causes intellectual disability, and dysmorphism (Emanuel syndrome) ^2^. In contrast, recombination in ova occurs in fetal life and then meiosis is arrested until puberty ^39^.

**Fig. 7.**
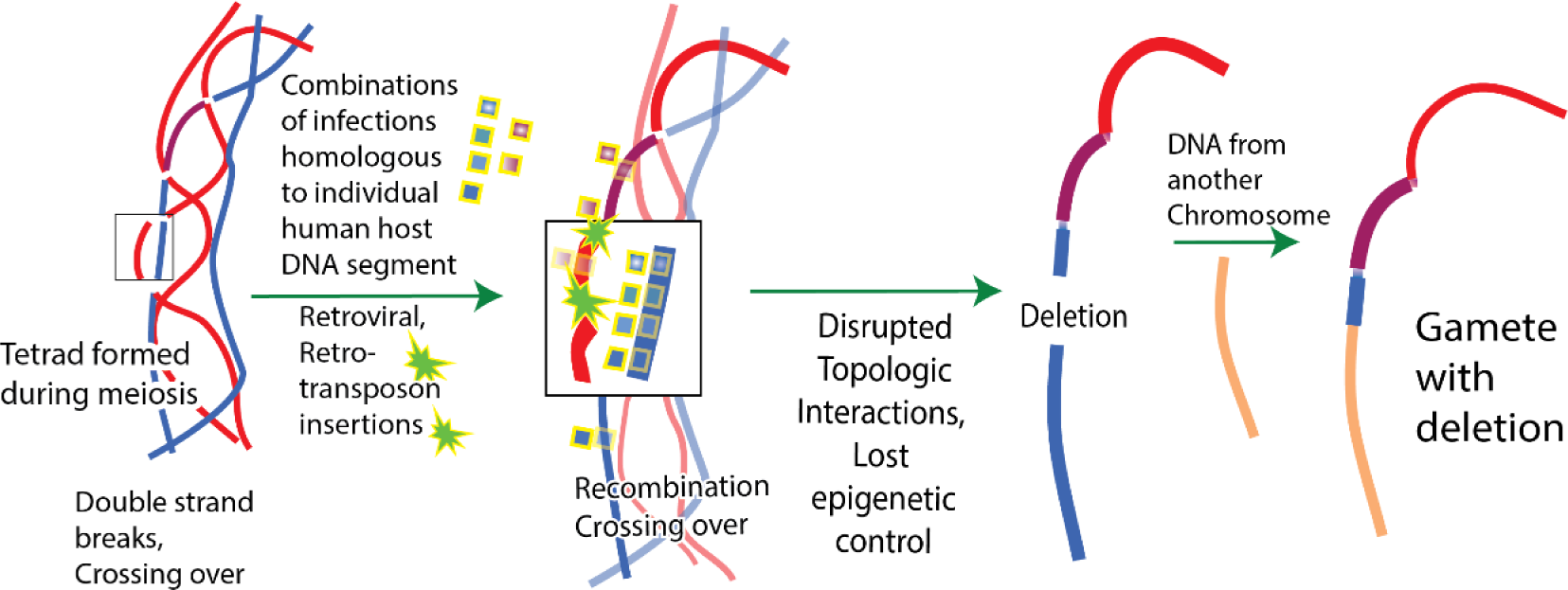
Model for interference with meiosis by DNA from infections leading to neurodevelopmental disorders. After duplication of parental chromosomes prior to generation of gametes, DNA from infections associates with strands of DNA in the many places including repetitive sequences of human DNA closely matching infection DNA. In addition, retroviruses and retrotransposons may integrate their DNA. Retroviral insertions are usually inactivated, but they can still interfere with normal processes such as reductive cell division, topological relationships among chromosomes, epigenetic regulation and high-fidelity break repair. Generally, interference from microorganisms may favor the illegitimate combinations due to palindromes reported by Mishra and co-workers ^2^. Hundreds of DNA breaks occur during meiosis and incorrect repair of these breaks is known to occur 1, causing chromosome anomalies such as deletions (shown). Clustered genes responsible for linked functions and epigenetic regulation in neurodevelopment are lost or displaced. Other chromosome segments with microbial homology do not contain identified genes but may be essential control regions, insulators, or essential for chromatin structures. There are about one million Alu repeats alone in the human genome and many match sets of infections in individual neurodevelopment disorders. More private matches also occur. When combined with effects from other microorganisms this can cause severe disruption of base pairing in gamete generation.

The resemblance of foreign microbial DNA to host background DNA may be a major factor in selecting infection and in human ability to clear the infection. Only one rare defective gamete is modeled in Fig. 7 but the male generates four gametes during meiosis beginning at puberty. Only one gamete survives in the female because three polar bodies are generated. Both balanced and unbalanced translocations can give rise to chromosome anomalies such as deletions and insertions because infection DNA can insert itself (e.g. exogenous or endogenous retroviruses) or interfere with break repairs during meiosis. A preexisting balanced chromosomal translocation in the family ^3^ increases the chances of generating a defective gamete during meiosis.

### Tests for artifacts in matches to human-microbial DNA

Genome rearrangements for patient data in Fig. 3 ^1^ produced 1986 matches with E<=e-10 and a mean value of 83% identity to microbial sequences (range 66-100%). About 190 Alu sequences resembled microbial sequences, supporting the idea that homologies among repetitive human sequences and microbes are real. Correspondence between microbial sequences and multiple human repetitive sequences increases possibilities that microbial sequences can interfere with essential human processes. Contamination of microbial DNA sequences by human Alu elements ^40^, was tested by comparing about 450 AluJ, AluS and AluY sequences to all viruses and bacteria in the NCBI database. In contrast, LINE-1 elements (NM_001330612, NM_001353293.1, NM_001353279.1) had no significant homology to bacteria and viruses.

The 1986 microbial homologues to human DNA near chromosome breakpoints in neurodevelopmental disorders were composed of 126 different viral or bacterial sequences. 24 different strains of Neisseria meningitidis in patient DGAP159 were 100% identical to sequences within human genomic rearrangements. (Total scores 385-637, E= 8e-13). The 25^th^ strain was 98% identical. Clostridium specifically matched sections of human ATPases, and ATP synthases, mitochondrial G elongation factor, and heat shock protein hsp70 mRNA. A 1004 bp fragment of waddlia chondrophila occurred 32 times near neurodevelopmental breakpoints with full-length 89% homology to human calcyclin binding protein (E=0.0). Waddlia chondrophila also had strong matches to human caspase recruitment. Another 55 of 126 viral / bacterial sequences were homologous to Alu sequences. Both non-Alu and Alu regions account for 90% (113 of 126) of the sequences homologous to microbes and present near breakpoints in human neurodevelopmental disorders.

### Tests for DNA sequence artifacts

To further test the possibility that some versions of these microbial sequences were sequencing artifacts or contaminated by human genomes, microbial genomes represented in the Figures and other related microbial genomes were tested for homology to human genomes.

Homologies to human sequences were found across multiple strains of the same microorganism (Table 3). In other cases, homology to some human gene product was found but there was not sufficient data available to compare multiple versions of the same microorganism. In these cases, the human gene was tested against all bacteria, and viruses. There were hundreds of significant matches. The complete clostridium botulinum genome had significant homology to human enzyme CDKL2 (86%) on chromosome 4. The region on chromosome 4: 579,587-75,632,298 spanned 52712 bp and was filled with over 50 Alu repeats, including multiple versions of AluS, AluY, and AluJ. Clostridium also had significant homology with long stretches on other chromosomes including chromosomes 2, 3, 10, 15, 1, 5 with 74% homology over about 2853 bp E=0.0. Similarly, the full genome of waddlia chondrophila (2.1 million bp) was homologous to multiple human genes (Table 3).

**Table 3.**
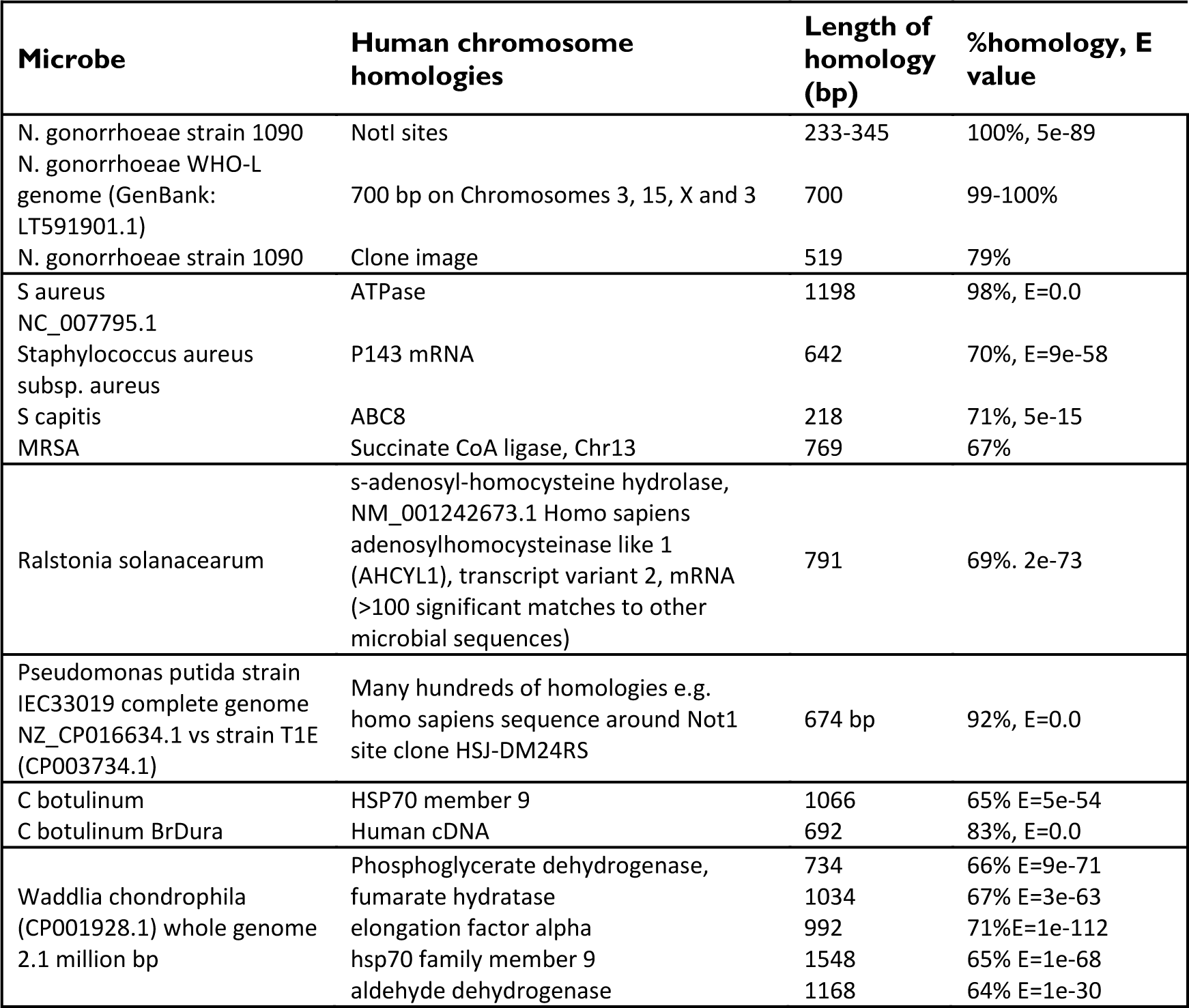
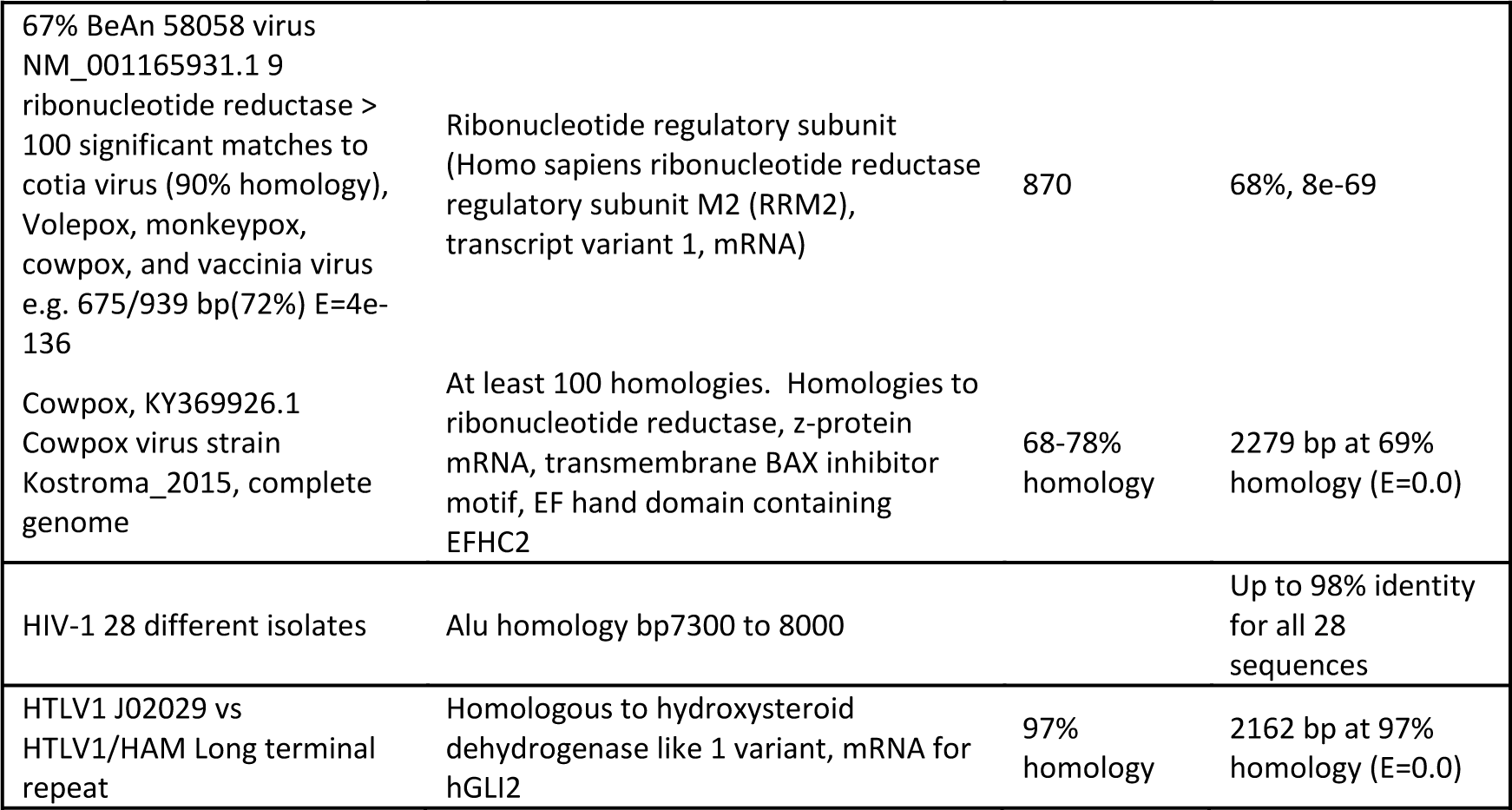
Independent evidence that microbial genomes have regions of homology to human DNA as predicted by results.

Retroviral infections are known to insert their sequences in human DNA, destroy cells, or have other methods of infection and transmission, such as by inhibiting key protective mechanisms. HIV-1 and other retroviral insertions matched congenital abnormalities in neurodevelopmental disorders. HIV-1 isolate SC007 (GenBank: KY713228.1) had 85% identity over 686 bases to human IgG2 lambda antibody and 82% homology to 412 bp of human DNA surrounding a rare NotI restriction site. To test for human sequence contaminants in these matching HIV sequences, Alu homologous regions of the HIV-1 genome (bp 7300-9000) in 28 different HIV-1 isolates were compared. All 28 HIV sequences matched the same region of human DNA, at up to 98% identity. The best match was to a LINCRNA on chr11:74,929,289-74,929,412. In contrast, only one zika virus sequence matched humans but 20 additional zika virus sequences did not, so zika virus was not considered further.

## Discussion

Long stretches of DNA in many infections match repetitive human DNA sequences which occur hundreds of thousands of times. Individual microorganisms also match non-repetitive sequences. Human infections may be selected for and initially tolerated because of these matches. It is almost impossible to completely exclude the possibility of sequencing artifacts or contamination of microbial sequences with human sequences. However rather than reflecting widespread, wholesale error due to human DNA contamination in many laboratories over many years, microbial homologies more likely suggest that infection sequences have been selected because they are homologous to regions of human DNA. This may be a driving force behind the much slower evolution of Alu DNA.

Infections can cause mutations by interfering with the most active period of recombination which occurs during the generation of gametes, an ongoing process in males. Infections such as exogenous or endogenous retroviruses are known to insert into DNA hotspots 38. Human DNA sequences that closely match infection DNA are proposed to generate chromosomal anomalies because similarities between microbial and human DNA interfere with meiosis (Fig. 7). Thus, the genetic background of an individual may be a key factor in determining the susceptibility to infection and to the effects of infection. At the genetic level, this suggests selective pressures for infections to develop and use genes that are similar to human genes and to silence or mutate genes that are immunogenic. Infection genomes evolve rapidly on transfer to a new host ^41^. Infections with long stretches of DNA nearly identical to human DNA makes the infection more difficult to recognize as non-self. For example, there is no state of immunity to *N. gonorrhoeae*. Long stretches of *N. gonorrhoeae* DNA are almost identical to human DNA. Alu sequences and other repetitive elements are thought to underlie some diseases by interfering with correct homologous recombination ^42^ or normal splicing. Infection DNA that is homologous to multiple, long stretches of human DNA may mask proper recombination or splice sites and encourage this abnormal behavior. Insertions of retroviral sequences are often epigenetically inactivated but can still interfere with meiosis and other normal processes (Fig. 7).

Neurons and cells in the immune system interact, sensing and adapting to their common environment. These interactions prevent multiple pathological changes in many disorders ^43^. Many genes that are implicated in neurodevelopmental diseases, reflect the strong relationship between the immune system and the nervous system. It was always possible to find functions within the immune system for genes involved in neurodevelopmental disorders (Figs. 1 and 4). Damage to genes essential to prevent infection leads to more global developmental neurologic defects including intellectual disability. These microbial homologies include known teratogens. Analyses of mutations within clusters of genes deleted in neurodevelopmental disorders predict loss of brain-circulatory barriers, facilitating infections. Damage to cellular genes essential for autophagy may lead to abnormal pruning of neural connections during postnatal development.

Aggregated gene damage accounts for immune, circulatory, and structural deficits that accompany neurologic deficits. Other gene losses listed in Table 1 and in deleted chromosome segments (Fig. 1) account for deficits in cardiac function, cell barriers, bone structure, skull size, muscle tone and many other non-neurologic signs of neurodevelopmental disorders.

The arrangement of genes in clusters converging on the same biological process may simplify the regulation and coordination between neurons and other genes during neurodevelopment and neuroplasticity. Genes that are required for related functions, requiring coordinated regulation have been shown to be organized into individual topologically associated domains 44. Functions that must be synchronized are grouped together on the same chromosome region and can be lost together by deletions (Fig.1) or by mutations of epigenetic controls (Table 1). Neurons are intimately connected to chromatin architecture and epigenetic controls ^45^. Homology to microbial infections anywhere in the cluster can then ruin complex coordinated neurological processes.

Microbial DNA sequences are unlikely to be contaminants or sequencing artifacts. They are all found connected to human DNA in disease chromosomes; multiple microbial sequences from different laboratories are all homologous to the same Alu sequence. Alu element-containing RNA polymerase II transcripts (AluRNAs) determine nucleolar structure and rRNA synthesis and may regulate nucleolar assembly as the cell cycle progresses and as the cell adapts to external signals ^46^. HIV-1 integration occurs with some preference near or within Alu repeats ^47^. Alu sequences are largely inactive retrotransposons, but some human-microbial homologies detected may be due to insertions from Alu or other repetitive sequences. Neuronal progenitors may support de novo retrotransposition in response to the environment or maternal factors ^48^.

Their variability and rarity make neurologic disorders difficult to study by conventional approaches. The techniques used here might eventually add to emerging disease prediction from medical imaging ^49^ and help predict lifelong susceptibility to a given infection. However, the methods used await new developments enabling the ability to distinguish infection by one microorganism from multiple microorganisms.

## Conclusions

1. DNAs in some congenital neurodevelopmental disorders closely matches multiple infections that extend over long linear stretches of human DNA and often involve repetitive human DNA sequences. The affected sequences are shown to exist as linear clusters of genes closely spaced in two dimensions.
2. Interference from infection can delete or damage human gene clusters and epigenetic regulators that coordinate neurodevelopment. This genomic interference accounts for immune, circulatory, and structural deficits that accompany neurologic deficits.
3. Neurodevelopmental disorders are proposed to begin when parental infections cause insertions or interfere with meiosis at both repetitive and unique human DNA sequences.
4. Congenital neurodevelopmental disorders are thus viewed as resulting from an assault on human DNA by microorganisms and an example of the selection of infecting microorganisms based on their similarity to host DNA.

## Acknowledgments

This work is based on the outstanding DNA sequence information and publications from investigators listed in the references. These results emphasize the importance of their work. It is a pleasure to acknowledge the inspiration, challenge and stimulation provided by the Department of Biochemistry and Molecular Genetics and the College of Medicine at UIC.

